# Spatial Variation in Cortex Glia Cell Cycle Supports Central Nervous System Organization in *Drosophila*

**DOI:** 10.64898/2026.01.05.697670

**Authors:** Vaishali Yadav, Syona Tiwari, Meenal Meshram, Ramkrishna Mishra, Rakesh Pandey, Richa Arya

## Abstract

Cortex glia (CG) in the ventral nerve cord (VNC) utilize mechanisms such as endocycling and acytokinetic mitosis to increase their nuclear count and support neighboring neural cells. However, the regulation of their cell cycle is not well understood. Our study examines larval development and reveals that CG nuclei in the thoracic region (tVNC) undergo successive rounds of replication and division, exponentially increasing their number to form a syncytial network around neural cells. In contrast, nuclei in the abdominal region (aVNC) only enhance their DNA content by endocycling. Key regulators, including cyclins and cyclin-dependent kinases, are vital for managing replication and division in CG. Notably, the M-phase regulator String (*stg*) is crucial for maintaining diploidy in tVNC nuclei and facilitating syncytial network formation. We demonstrate that ectopic *stg* expression can induce nuclear division in hyperploid aVNC CG. This study highlights regulatory mechanisms in CG, emphasizing the importance of region-specific cell cycle regulation for nervous system organization.

## Introduction

The central nervous system (CNS) is a complex network comprising various cell types, including neurons, glia, and neuroblasts, which are also known as neural stem cells (NSCs). Each of these cell types plays essential role in the organization and function of the CNS. In the *Drosophila* CNS, multiple glial subtypes have been classified based on their morphology. These include surface glia (perineural and sub-perineural glia), cortex glia (CG), and neuropil-associated glia (wrapping glia and astrocyte glia)(Freeman, 2015; Ito et al., 1995; Stork et al., 2012).

Sub-perineural and CG cells increase their DNA content and become polyploid (Unhavaithaya and Orr-Weaver, 2012; Von Stetina et al., 2018; Rujano et al., 2022). These cell types form a syncytium through both common and distinct cell cycle pathways. Sub-perineural glial cells, which create the functional blood-brain barrier, undergo polyploidization through endocycling, after which their nuclei divide via endomitosis (Unhavaithaya & Orr-Weaver, 2012; Von Stetina et al., 2018) Similarly, CG cells also reportedly become polyploid by increasing their DNA content through endocycling and form syncytia through endomitosis and acytokinetic mitosis (Rujano et al., 2022).

Polyploidy can be observed in various cell types across the animal kingdom. In *Drosophila*, several larval tissues—including the salivary gland, ovary/follicle cells, hindgut/rectal papillae, and brain/subperineural glia—are known to attain a hyperploid state during normal development. This increased ploidy allows them to secrete specialized proteins to perform unique functions (Hammond and Laird 1985; Spradling A, Mahowald A 1980; Fox &Duronio, 2013;Unhavaithaya & Orr-Weaver, 2012; Von Stetina et al., 2018) In mammals, placental trophoblast giant cells (TGCs), keratinocytes, hepatocytes, and blood megakaryocytes (MgKs) also exhibit polyploidy (Barlow & Sherman, 1972; Brown et al., 1997; Kudryavtsev et al., 1993; Lordier et al., 2008; MacAuley et al., 1998; Odell et al., 1976). Megakaryocytes undergo endomitosis as part of the platelet production process (Ravid et al., 2002) while TGCs enter the endocycle to act as a barrier between maternal and embryonic tissues (Rossant& Cross, 2001; Watson & Cross, 2005). Polyploidy is also recognized in bacteria (Mendell et al., 2008) and in other multicellular organisms, such as ciliated protozoa (Yin et al., 2010). Increased ploidy can be achieved through various mechanisms, including endocycle, cell fusion, endomitosis, acytokinetic mitosis, chromosome segregation failures, and impaired mitosis in checkpoint-defective cells (Britton & Edgar, 1998; Edgar & Orr-Weaver, 2001; Lilly &Spradling, 1996; Shabo et al., 2020; Ullah et al., 2009; Unhavaithaya & Orr-Weaver, 2012b; Wong & Stearns, 2005; Zanet et al., 2010).

CG is one of the major types of glial cells born during the mid-embryonic stage (Pereanu et al. 2005; Coutinho-Budd et al. 2017; Rujano et al. 2022). Recently, it has been suggested that during the early larval period, CG cells grow and increase their DNA content through endocycling. Later, their nuclei multiply via endomitosis and acytokinetic mitosis, without undergoing traditional cell division (Rujano et al., 2022). These CG cells form a complex 3D reticulate structure through homotypic cell–cell fusion, resulting in an extensively connected cytoplasmic niche around neural cells that supports their placement, trophic needs, and proliferation (Banach-Latapy et al., 2023; Coutinho-Budd et al., 2017; Dumstrei et al., 2003; Plazaola Sasieta et al., 2019; Read, 2018; Rujano et al., 2022; Spéder& Brand, 2018). Additionally, CG cells also protect NSCs from damage caused by oxidative stress and nutrient deficiencies (Bailey et al., 2015; Cheng et al., 2011). Adhesion of the CG niche non-autonomously regulates the fate of NSCs (Cristina et al., 2024). Previously, we demonstrated that the transcription factor Cut influences CG development by regulating ploidy and the formation of complex cellular architecture around neural cells (Yadav et al., 2024). The absence of Cut disrupts DNA replication, preventing CG nuclei from increasing their DNA content, ultimately leading to their removal from the nervous system (Yadav et al., 2024). However, the mechanism by which CG with a complex reticulate architecture expands within the continuously developing CNS remains unresolved. While it is suggested that these cells form syncytia through acytokinetic mitosis, the role of cell cycle regulators in this process is debated. Surprisingly, some key cell cycle regulators, such as dacapo (*dap*) for S phase and String (stg) for M phase progression, have been shown to play no role in these cells (Rujano et al., 2022). In contrast, another study suggests that stg and *CycA* influence nuclear division without affecting the complex architecture of CG (Beachum et al., 2025).

Here, we conducted a comprehensive examination of the functions of various cell cycle regulators in CG growth. We evaluated the roles of genes regulating different phases of the cell cycle, including G1, S, G2, and M. Interestingly, our study reveals that most key cell cycle regulators—except for DNA repair genes—are necessary for the regulation of CG cell cycle and overall cellular growth, highlighting the intricate relationship between cell cycle control and glial cell development. We observed that CG in different regions of the ventral nerve cord (VNC), including the thoracic (tVNC) and ventral (aVNC)regions, displays varying levels of complexity. In the tVNC, CG undergo multiple rounds of replication and nuclear divisions in an exponential manner forming a syncytium of largely diploid nuclei. In contrast, the CG in aVNC region do not make syncytia, instead undergo endocycle to increase their DNA content. These cells increase their DNA content slowly in a linear fashion. Collectively, our findings show that while CG follow a variant cell cycle for their growth, canonical cell cycle regulators play a pivotal role in their development. Furthermore, CG development is not uniform throughout the CNS; it adapts to the specific needs and functions of different regions of the CNS, potentially contributing to the functional diversity observed across various areas of the nervous system.

## Results

### Cortex glia nuclei in tVNC undergo multiple rounds of replication and nuclear division during larval development

The CG in tVNC displays a complex, dynamic pattern of nuclear division and DNA replication during larval development, including endomitosis, acytokinetic mitosis, and endocycling (Rujano et al., 2022). Since CG cells fuse to form syncytial units, it is difficult to precisely count their number or determine their ploidy at different larval developmental time points. To understand the mechanisms of nuclear division and CG growth, we counted the number of nuclei and the DNA content per nucleus across different larval stages in tVNC (T1-A2)We collected data at the following developmental time points: ALH 0h, 24h, 40h, 48h, 65 h 72h, 90hr and 96h (Fig. 1A-F).

**Fig 1:**
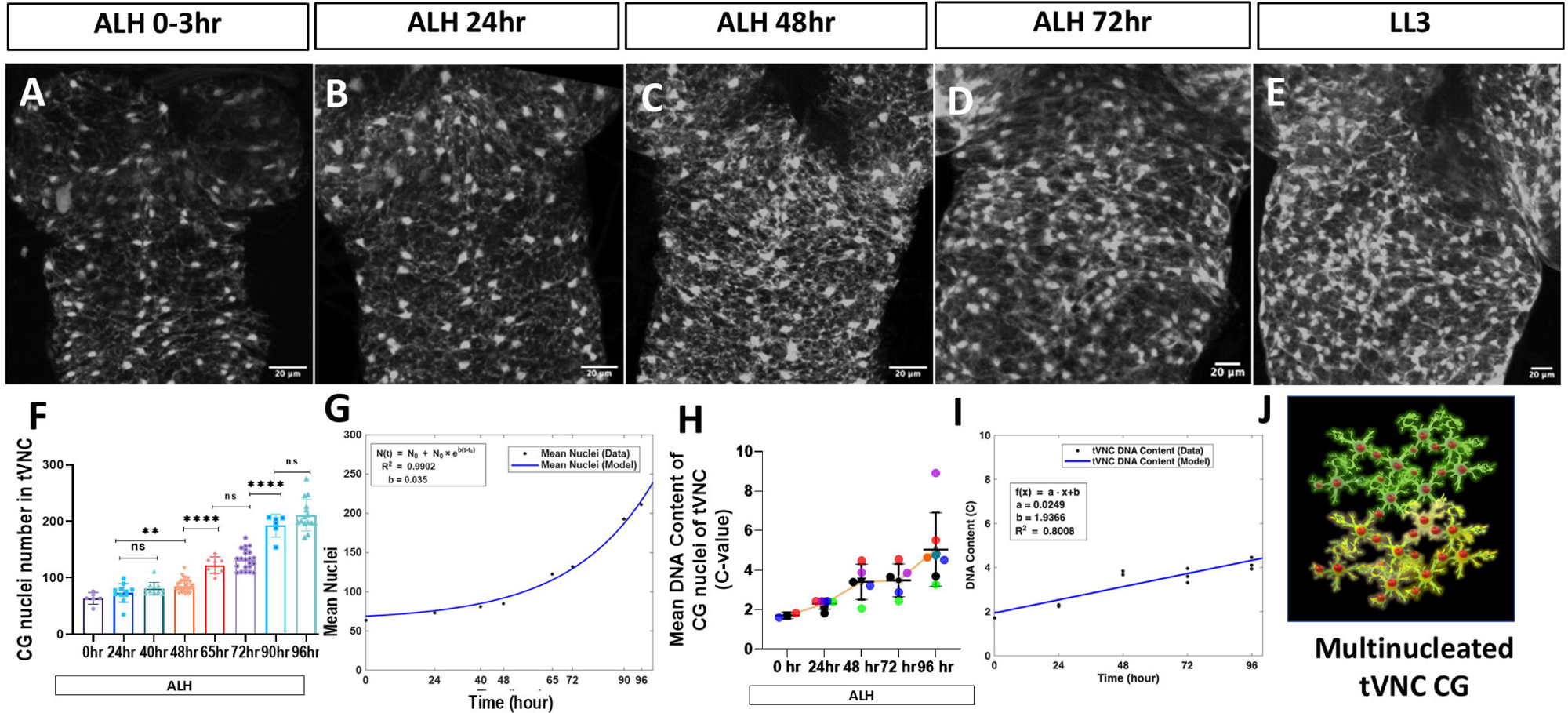
Cortex glia achieves hyperploidy by undergoing multiple rounds of replication and nuclear division during larval development. A-E) Evaluation of CG nuclei and network organization during larval development in the tVNC at ALH 0 hr, ALH 24 hr, ALH 48 hr, ALH72 hr, and ALH96 hr (at 25 °C) F, H) Quantification of CG nuclei number and DNA content with respect to sperm in tVNC at different developmental stages from ALH 0hr to ALH96 hr (at 25 °C) respectively. The colored dots represent the number of sets analyzed. G) Mathematical model showing that CG nuclei number increases exponentially during the larval developmental stages from ALH0 to ALH96 hr I) Mathematical model showing that CG mean DNA content increases linearly during the larval developmental stages from ALH0 to ALH96 hr J) Model showing two tVNC multinucleated CG in green and yellow are connected. The statistical evaluation of significance based on unpaired *t*-test is marked with asterisks, \*\*\*\**P* < 0.0001, \*\*\**P* < 0.001, and \*\**P* < 0.01. Scale bar: 20 µm

The number of CG nuclei remain almost unchanged between ALH 0hr to 40h, indicating no nuclear division has happened (Fig. 1A-F) (Coutinho-Budd et al., 2017). However, from ALH 48h onward, the number of nuclei starts increasing, more prominently between ALH 72-96, indicating the occurrence of rapid nuclear division (Fig. 1F,H)(Coutinho-Budd et al., 2017). To understand the rate of replication in individual CG nuclei, we measured their DNA content during each developmental time point and normalized it with sperm DNA (details in the methods section). We noted that from ALH 24h onwards, the DNA content per CG nucleus is consistently greater than 2C, indicating a hyperdiploid state. In fact, during ALH 0-40h, the mean DNA content per nucleus increases to 3.43C (Fig 1H); but the number of nuclei does not increase drastically, indicating an ongoing replication (Fig. 1F,G). Between ALH 48-72h, almost half of the CG nuclei were divided, and thus the mean number of nuclei increased from 85 to 137 (Fig 1F,H).

To understand the rate of increase in nuclear number, we fitted the data to a mathematical model N(t)=No+exp(b(t-t_o_) (method section Eq. 1) using MATLAB. Where N(t) is the mean nuclei number at any time t, N_0_ is the initial mean nuclei number, b represents the rate of increase in nuclei number, and t_0_ indicates the time after which a rapid rise in nuclei number is observed. We observed that the model represented by Eq. 1 fits the observed data very well with R square value=0.98. Hence, with both the observed and modelled data, we infer that the mean number of nuclei increases very slowly till t_o_=65 h and increases exponentially after that (Fig 1G).

Interestingly, despite nuclear division, the DNA content per nucleus remained largely within the range of 2C to 4C from ALH 0 h to ALH 72 h, indicating that both nuclear division and replication occur during these stages (Fig 1F-H). Similarly, from ALH 72h to 96h, the mean nuclei number further increased from 132 to 211, showing another round of nuclear division. During this window of development, the DNA content per nucleus increased, ranging from 3C to 5C, indicating continued replication alongside nuclear division (Fig 1F-H).

Further to understand the rate of change in DNA content, we modelled the data using the quartiles Q1, Q2 and Q3 of CG to show heterogenous population. Here we found that quartiles DNA of tVNC shows a linear pattern across the developmental stages (Fig. 1I). These observations altogether indicate that during ALH 0-40h, the CG nuclei increase their DNA content up to 4C without dividing. In contrast, during ALH 48-70h, these nuclei begin to divide while maintaining a DNA content between 2C and 4C. However, in the later stages of larval development (LL3), the DNA content per nucleus increases beyond 4C, suggesting entry of a population of nuclei into an endoreplication mode of replication. Therefore, while the CG nuclei in the tVNC region are primarily diploid and complete two round of nuclear divisions, they increase their DNA content by endocycling in the later larval stages. Since tVNC cortex glia are syncytial, we can conclude that these cells are polyploid, while the individual nuclei are largely diploid.

### 2. *Cyclin E* and *dacapo* are necessary and sufficient for the regulation of tVNC cortex glia cell cycle

The dual function of *CycE/Cdk2* is crucial for maintaining genomic stability and ensuring proper cell cycle progression (Thomer et al., 2004). On one hand, *CycE/Cdk2* phosphorylates the pre-replication complex, leading to its inactivation; on the other hand, it serves as an entry point into S-phase, initiating DNA synthesis (Calvi et al., 1998; Narbonne-Reveau& Lilly, 2009; Thomer et al., 2004; Whittaker et al., 2000).

We observed that knocking down *CycE* significantly hinders the development of CG in tVNC region (see Fig. 2A-C’, G*; CycE-RNAi*#2; Fig. S2B*CycE-RNAi* #1). During ALH 24h and ALH 48h, the number of CG nuclei remains constant in both control and *CycE* knockdown conditions. However, from ALH 72h to LL3, while the number of nuclei in the control increases, it remains relatively stable in the *CycE* knockdown group, keeping the number similar to that of ALH48h (see Fig. 2E).

**Fig 2:**
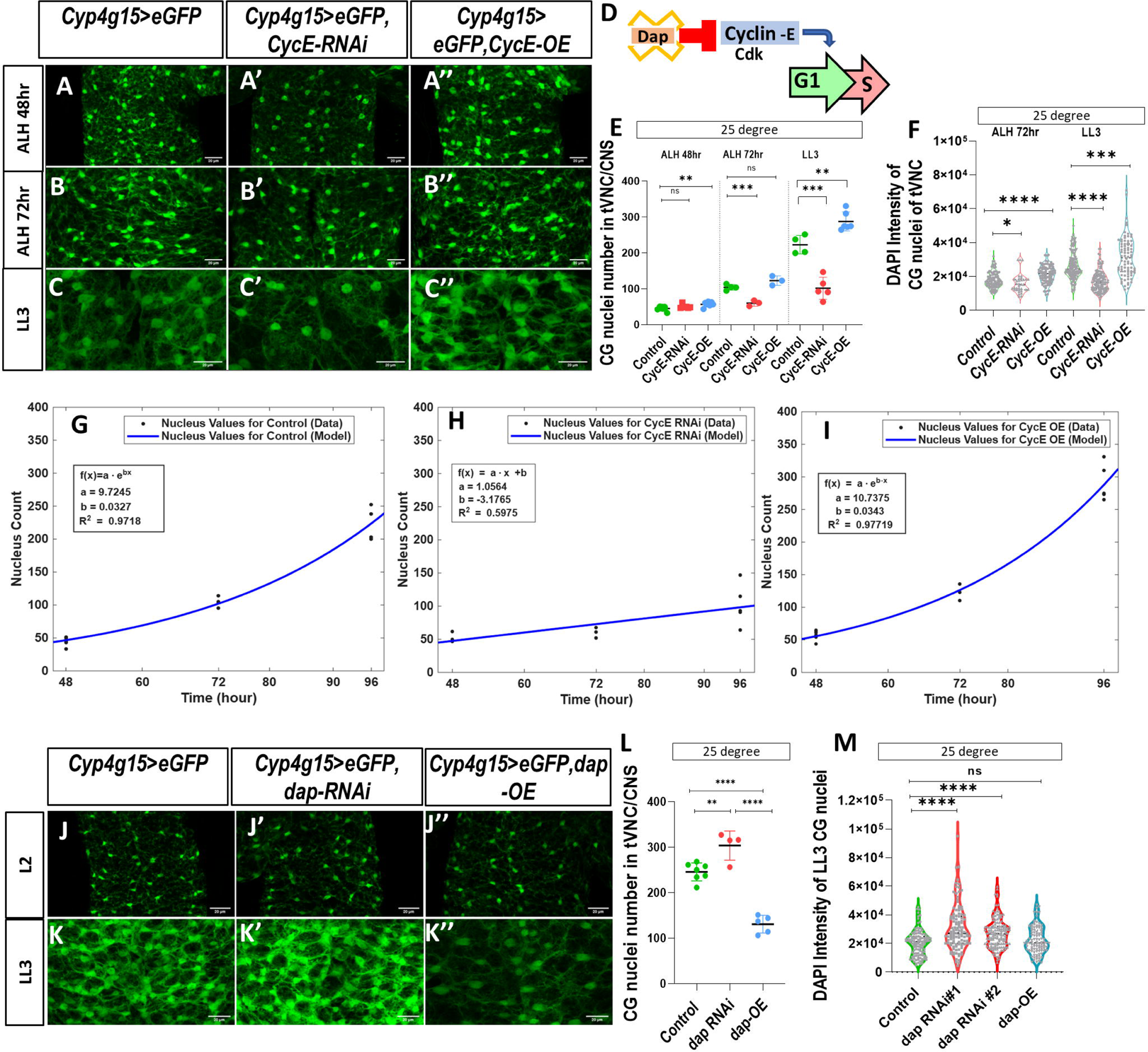
*Cyc-E* & *dacapo* regulate the G1 to S-phase transition of the tVNC Cortex glia cell cycle. A-C’’) Ventral sections of confocal projections, A, B, C) Control (*Cyp4g15>eGFP*) having3-4 CG nuclei per hemisegment, making a patterned arrangement as in embryonic life. A’) *CycE* defective CG (*Cyp4g15>eGFP; CycE-RNAi*#2) shows no significant change in nuclei number at ALH 48 hr B’, C’) while there is significant reduction at ALH 72 hr and LL3 stages. Upon ectopic expression of *CycE* (*Cyp4G15>eGFP; CycE-OE#1*), nuclei number increases at A’’) ALH 48hr, B’’) ALH72hr, and C’’) LL3 stage. D) A schematic diagram shows the regulation of G1-S transition by *CycE-CDK* complex, where *dacapo* is shown as an inhibitor of the complex. E) Quantification of CG nuclei number in tVNC upon misexpression of *CycE* F,) DNA content in CG in tVNC upon misexpression of *CycE* G, H, I) Mathematical model showing that tVNC CG nuclei number increases exponentially in *CycEOE* with almost the same rate as in control (G, J). While in *CycE-RNAi*, the number of nuclei increases linearly (H). J-J”) At the L2 stage, misexpressing da*capo* does not change the nuclei number (J’-J”), K-K”’) LL3 stage, there is a significant increase in the number of nuclei in dap knockdown (*Cyp4g15>eGFP;dap-RNAi*#1), while overexpression of dap (*Cyp4g15>eGFP;dap-OE*#1) shows less increase in nuclei compared to control. L) Quantification of CG nuclei number in tVNC upon misexpression of *dap*. M) DNA content in CG in tVNC upon misexpression of *dap*. The statistical evaluation of significance based on unpaired *t*-test is marked with asterisks, \*\*\*\**P* < 0.0001, \*\*\**P* < 0.001, and \*\**P* < 0.01. Scale bar: 20 µm J-J”) At L2 stage, misexpressing *dacapo* does not change the nuclei number in (J’-J”) and in K-K”’) LL3 stage there is a significant increase in the number of nuclei in *dap* knockdown K’) knockdown condition (*Cyp4g15>eGFP;*d*ap-RNAi*#1) while in K’’) overexpression (*Cyp4g15>eGFP;*dap*-OE*#1) condition the nuclei number is less in comparison to control. The statistical evaluation of significance based on unpaired *t*-test is marked with asterisks, \*\*\*\**P* < 0.0001, \*\*\**P* < 0.001, and \*\**P* < 0.01. Scale bar: 20 µm

Using mathematical modeling, we determined that under normal conditions, the number of CG nuclei increases exponentially from 48h to LL3 (exponential growth rate, b = 0.0327) (Fig 2G). In contrast, with *CycE* knockdown, the rate of nuclear increase is constant (slope of line, a = 1.05) and follows a linear pattern (Fig.2H). Conversely, upon *CycE* overexpression, the number of nuclei continues to rise with a higher exponential rate (b = 0.0343) (see Fig. 2A-C’’, E, I; *CycE-OE*#1; Fig. S2B*CycE-OE*#2).

Since *CycE* is essential for DNA replication, we measured DAPI content and found that the nuclei of CG with *CycE* knockdown had significantly less DNA than those in the control at ALH 72h and LL3, indicating delayed replication and impaired nuclear division (Fig. 2F, H). Interestingly, the DAPI intensity in *CycE*-overexpressing CG nuclei was higher than that in the control at ALH 72h and LL3, suggesting that these nuclei enter the endocycling phase earlier than the controls (Fig. 2F, I).

In *Drosophila*, *CycE/Cdk* levels are regulated by *dacapo* (*dap*), a member of the CIK/KIP family which binds and inhibits CycE-Cdk2 activity (Audibert et al., 2005; De Nooij et al., 1996; Lane et al., 1996). Upregulation of *dap* is necessary after the last mitosis to arrest cells in G1/G0 before terminal differentiation in many post-mitotic cell types (De Nooij et al., 1996). To determine if *dap* is important in the CG cell cycle, we modulated its levels by its knock down and overexpression. Knockdown of *dap* in CG results in an increase in the number of nuclei (Fig. 2, J-K’’, *dap-RNAi* 36720, Fig. S2C*dap-RNAi* 64026). This phenotype resembles *CycE* overexpression in CG, having a higher rate of replication. On the other hand, dap overexpressing CG does not significantly increase the number of their nuclei, as seen in the case of *CycE* knockdown (Fig. 2K”). Overall, we found that *CycE* and its negative regulator dap are essential for replication in CG.

### 3. M-phase regulator *string* and *cdk1* are required for nuclear division in the tVNCcortex glia

The CG cells in tVNC undergo a variant cell cycle to form syncytial units with varying DNA content in individual nuclei (Rujano et al., 2022; Yadav et al., 2024). It’s important to note that String (stg), a tyrosine-protein phosphatase, activates the mitotic cyclin/Cdk complex necessary for a cell’s entry into mitosis. In contrast, stg remains repressed in cells undergoing the endocycle(Deng et al., 2001; Di Talia &Wieschaus, 2012; Edgar et al., 1994; Edgar & O’Farrell, 1990). Although standard mitotic stages are observed during nuclear division in tVNC CG, the precise role of stg remains unclear(Rujano et al., 2022).

Given that CG glia in the tVNC region increase their nuclear number exponentially (Fig. 1H, I), we investigated whether stg influences CG cell-cycle regulation. In *stg*-depleted CG cells, the number of nuclei did not increase from ALH48 (mid-L3) onwards, indicating a failure in nuclear division (Fig. 3A’, B). Interestingly, the DNA content per nucleus in *stg*-depleted CG nuclei increased significantly reaching 10C-14Cat ALH 96hr, suggesting that DNA replication continued uninterrupted in these nuclei even in absence of nuclear division (Fig. 3F). This indicates that the loss of *stg* inhibits mitotic division in these CG cells, while S phase continues, leading to an increase in DNA content per nucleus.

**Fig 3:**
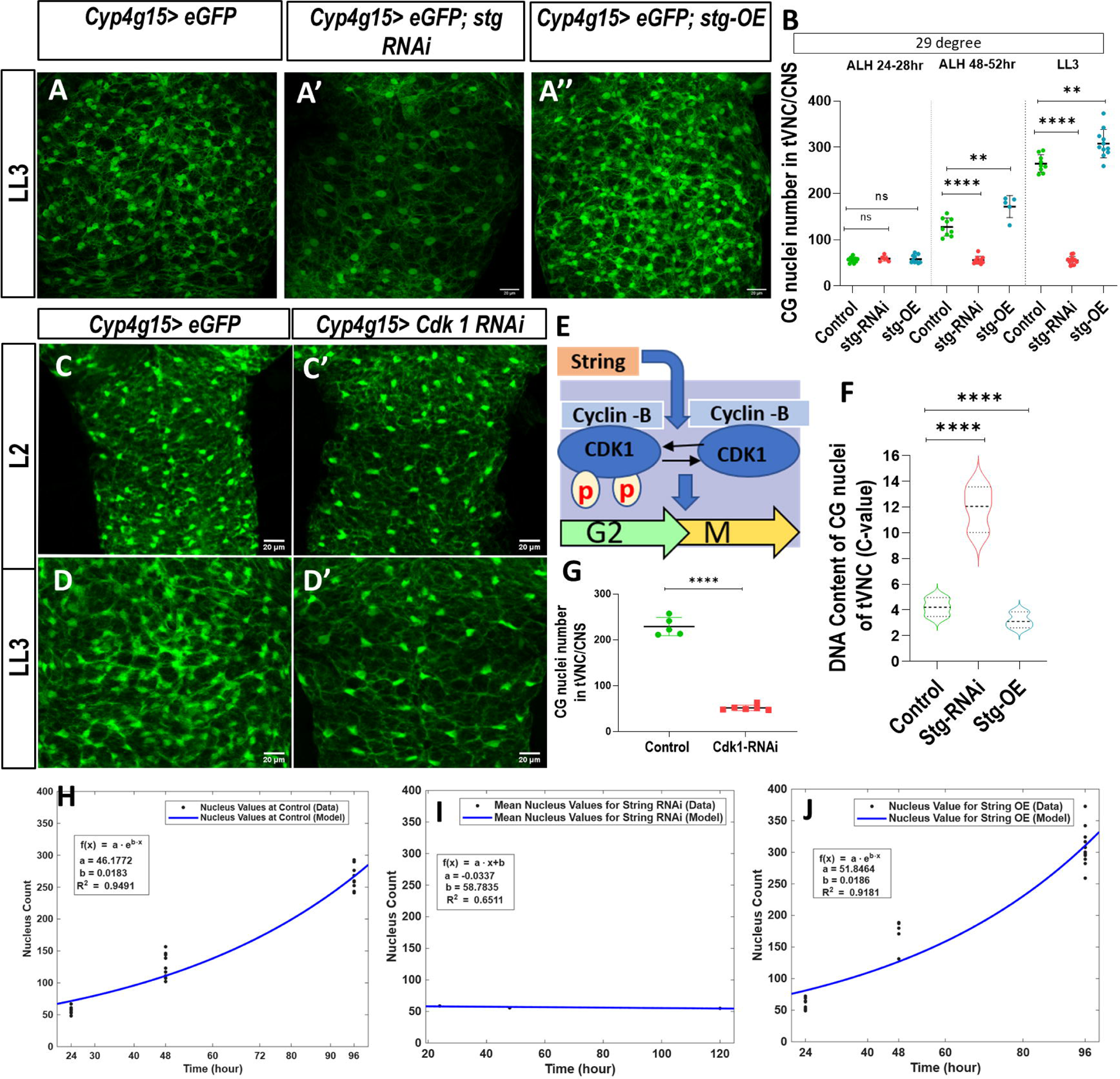
M-phase regulator *string* and *cdk1* are required for nuclear division in the tVNC cortex glia. A-A’’) *stg* defective CG (*Cyp4g15>eGFP; stg-RNAi*#3) shows fewer CG nuclei, while A”) ectopic expression of stg (*Cyp4g15>eGFP*; *stg-OE*) causes more nuclear division, resulting in more CG nuclei compare to control (A). B) Quantification of CG nuclei in *stg-RNAi* and *OE*. C, C’) Knockdown of *Cdk1* (*Cyp4g15>eGFP; cdk1-RNAi*#2), causes no significant change in (C’) CG nuclei number in L2 stage, while in LL3 stage D, D’) there are less number of nuclei, indicating that Cdk1 is also required for nuclear division. E) Model showing the *stg* removes the inhibitory phosphatase, thus activating the mitosis progression through *CycB/cdk* complex, F) DNA content (C-value) of CG nuclei showing that knockdown of *stg* significantly increases the DNA content, while its overexpression causes significant reduction in DNA content. G) Knockdown of *Cdk1* causes significant change in tVNC CG nuclei number. H-J) Mathematical model showing that tVNC CG nuclei number increases exponentially in *stg-OE* (J) with almost the same rate as in control (H). While in I) *stg-RNAi* condition, the change in nuclei number follow linear pattern. The statistical evaluation of significance based on unpaired *t*-test is marked with asterisks, \*\*\*\**P* < 0.0001, \*\*\**P* < 0.001, and \*\**P* < 0.01. Scale bar: 20 µm

Conversely, overexpression of *stg* resulted in an increased number of CG nuclei at mid-L3 and LL3 stages compared to the control (Fig. 3B). Notably, during the LL3 stage, the mean DNA content in the control varied between 2C and 5C per nucleus. However, *stg* overexpression reduced this and bring it to2C and 4C by inducing more nuclear divisions (Fig. 3F).

Furthermore, through mathematical modeling, we demonstrated that, unlike the control group, in which the number of CG nuclei increases exponentially from 48 hours to LL3 (b = 0.0183), the knockdown of stg reduces the rate of change and levels off, yielding a constant slope of −0.0337. This indicates a linear pattern (see Fig. 3H, I). In contrast, the overexpression of stg leads to an exponential increase in the number of nuclei, with an exponential rate of 0.0186 (represented by ‘b’) as shown in Fig. 3J.

We also checked another M-phase regulator, *Cdk1*, which is a member of the *CycB/Cdk* complex that regulates the cell cycle in tVNC CG cells. Knocking down *Cdk1* in CG cells interfered with nuclear division, preventing an increase in their number (Fig. 3C-D’, G). Therefore, we conclude that reducing both *stg* and mitotic *Cdk1* prevents CG from increasing their nuclear number.

### 4: Replication checkpoints are dispensable during tVNC cortex glia growth and nuclear division

To ensure accurate replication and division of cells, various checkpoints are present to maintain the integrity of the cell cycle (Kastan & Bartek, n.d.; Machida & Dutta, 2005). Checkpoints are largely known for their role in cell cycle progression, origin firing, and the stabilization of stalled replication forks (Nurse, 1994). In the thoracic CG, we examined the roles of the ATR/Chk1 and ATM/Chk2 replication checkpoints. Chk1 works as downstream effector kinase of ATR, and Chk2 acts as a downstream effector kinase of ATM (Smith et al, 2010). Surprisingly, we did not find any observable change in the number of nuclei upon knockdown of several replication checkpoints such as *grp*, *loki*, *tefu*, *ATRIP*, and *Chk1*(Fig. 4A-A’’,B). There was no-significant change in mean DNA content in *Chk1* knockdown condition (Fig. 4A’) Further, to check if ectopic expression of checkpoints could alter CG cell cycle, we ectopically express *Chk1*. Interestingly, we note that the ectopic presence of this checkpoint slows down the nuclear division, resulting in a 0.50-fold reduction in CG nuclei number at the LL3 stage (Fig. 4A-A’’, B). while there was a 0.28-fold increase in mean DNA intensity (Fig. 4C).

**Fig 4:**
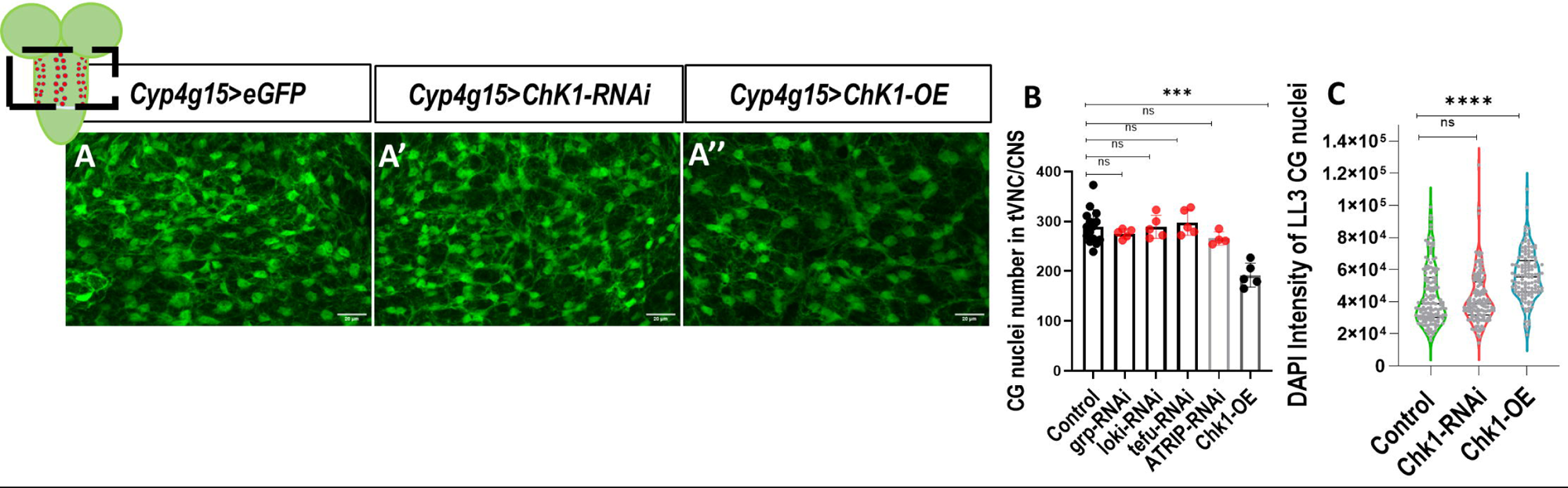
Replication checkpoints are dispensable during tVNC cortex glia growth and nuclear division. A’) Knockdown of *Chk1* in tVNC CG does not change the nuclei number while in A’’) *Chk1-OE* the number of nuclei is significantly less indicating that more of *Chk1* is inhibiting nuclear division in comparison to A) control. B) Quantification of CG nuclei number in tVNC/CNS in knockdown condition of grp (Chk1), loki (Chk2), tefu (ATM), ATRIP C) Quantification of DAPI intensity of tVNC CG The statistical evaluation of significance based on unpaired *t*-test is marked with asterisks, \*\*\*\**P* < 0.0001, \*\*\**P* < 0.001, and \*\**P* < 0.01. Scale bar: 20 µm

Since CG are a special type of cells following normal nuclear division as well as endocycling, we assume that the absence of replication checkpoints may allow them to bypass the typical cell cycle control which may interfere in achieving the hyperploid DNA state. Since these cells form syncytium and act as glial units, they may not be undergoing normal DNA replication or normal chromosome segregation. An abnormal distribution of chromosomes 2-3 in individual nuclei is also reported in these cells (Rujano et al., 2022). This adaptation, lacking strict checkpoint control, may allow these cells to switch between endocycle and nuclear division whenever needed. This unique cell-cycle regulation in tVNC CG highlights specialized adaptations that meet specific functional requirements.

### 5: CG in aVNC region endocycle without undergoing nuclear division

CG in aVNC exhibits a markedly different developmental pattern than the CG in tVNC. During larval development, these cells do not show an increase in the number of nuclei; rather, some of them are eliminated by an unknown mechanism (Fig. 5C). Using the Fly-FUCCI tool, we found that at ALH 24h, 92% of abdominal CG are in G1, 0.56% in S, and 6.52% in G2/M. At ALH 48h, 88% of CG are in G2/M phase, and thereafter, most of them remain in the G2 stage of the cell cycle (Fig. 5A, B).

**Fig 5:**
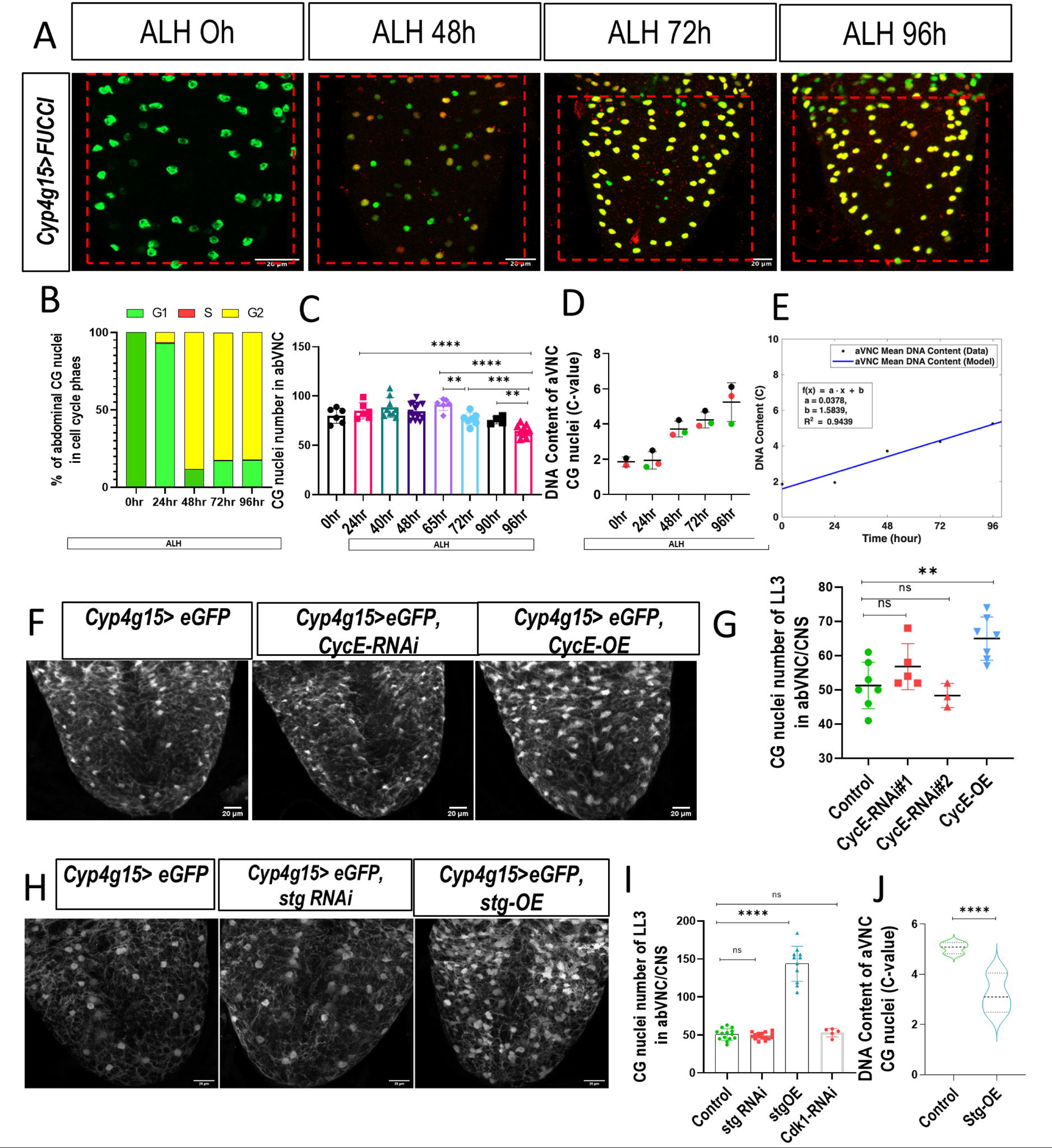
CG in aVNC region endocycle without undergoing nuclear division. A) G1 (E2F1, green), S (red) and G2/M (CycB, yellow) phases of the CG cell cycle detected with Fly-FUCCI at ALH 0hr, ALH 48hr, ALH72hr, ALH 96hr. B) Quantification of percentage of aVNC CG nuclei in different phases of cell cycle through fly-FUCCI. C) Quantification of CG nuclei number in aVNC at different developmental stages ALH 0hr, ALH 24hr, ALH 40hr, ALH 48hr, ALH 65hr, ALH 72hr, ALH 90hr, ALH 96hr. D) DNA content (C-value) of CG nuclei of aVNC, shows that content keep on increasing from ALH 0hr to ALH 96hr E) Mathematical modeling of DNA content shows a linear pattern of increase F) Knockdown of *CycE-RNAi*#1 and #2 does not change the number of CG nuclei, whereas CycE-OE forces them to enter mitosis and increases the number. G) Quantification of CG nuclei number in aVNC /CNS in *CycE-RNAi* and *OE* VNCs. H) knockdown of mitotic regulators like *stg* and *Cdk1* does not change the number of aVNC CG nuclei, while overexpressing *stg* makes them forces them to enter mitosis and increases the number. I) Quantification of CG nuclei number in aVNC /CNS in *stg-RNAi*#1, *Cdk1-RNAi*, *stg-OE#1* condition. J) DNA content (C-value) shows that in *stg* overexpression induces the nuclear division, bringing the DNA content down to 2-4C compared to control values, which are above 4C. The statistical evaluation of significance based on unpaired *t*-test is marked with asterisks, \*\*\*\**P* < 0.0001, \*\*\**P* < 0.001, and \*\**P* < 0.01. Scale bar: 20 µm

Interestingly, the analysis of DNA content in abdominal CG reveals a unique ploidy pattern. From ALH 0-48 hours, these cells maintain a consistent ploidy around 2C-4C, similar to thoracic CG (Fig. 5D). From ALH 48hr, the DNA content increases from 4C to 6C. Thus, abdominal CG undergo one complete and another ongoing round of replication until the LL3 stage (Fig. 5D, E). Mathematical modeling shows that abdominal CG cells increase their DNA content from ALH 0 to 96 hr in a linear manner (slope of the line is 0.0378) (Fig. 5E).

Furthermore, we assessed the effects of representative cell cycle regulators on abdominal CGs at the LL3 stage. Knockdown of *CycE*, *stg*, and *Cdk1* does not significantly change the number of CG nuclei (Fig. 5G,I). The ectopic expression of both *CycE* and *stg* forced these CG nuclei to divide (Fig. 5H, J). The total number of nuclei in the central abdomen (A1-A7) increased significantly upon *CycE* and *stg* overexpression (Fig. 5F-I). It is interesting to note that the DNA content in the *stg* overexpression condition was reduced to 2-4C due to induced nuclear division (Fig.5KJ), indicating that ectopic *stg* can force CG to divide and maintain a diploid status instead of undergoing endocycle. Together, this data indicates that CG in aVNC are competent to divide but remain arrested in G2 during normal development.

### 6. Region-specific differential regulation of the cell cycle determines the ploidy of CG nuclei

There is a distinct difference in the replication patterns of CG in tVNC and aVNC during the development of the larval CNS. The CG nuclei in tVNC divide and increase in number exponentially while maintaining their ploidy between 2C and 4C in most of the larval development and exceed beyond 4C in later larval life (Fig. 1 F, G, H). In contrast, the CG nuclei in the aVNC region do not divide; instead, they endocycle to increase their ploidy to 6C (Fig 5C-F).

We noted that *stg* knockdown completely stops nuclear division in tVNC CG, converting a normally dividing nucleus into an endocycling one, resulting in increased DNA ploidy, with a mean of 12C and a maximum of 14Cat ALH 96 hr (Fig. 5F).To understand how nuclear division might influence DNA content per nucleus in CG of tVNC where replication and nuclear division proceed simultaneously, we estimated the expected DNA content per nucleus by assuming a scenario in which CG nuclei in tVNC undergo only replication rounds without dividing using Equation 2. Since the initial number of nuclei in the CG of tVNC was around 64 during ALH 0h, we assume that under “no nuclear division” assumption, this nuclei number will remain constant throughout the larval development (Fig. 1F). This ‘no nuclear division’ assumption showed that DNA content in individual CG nuclei could reach upto20C if the nuclei fail to divide (Fig. 6A). The analysis suggests that during the larval life, one CG nucleus finish three complete replications, one from ALH 0h-35h (reaching to 4C), the second from ALH 48h-72h ( reaching to 8C), and the third from ALH 72h-90h (16C), and a fourth round starting from ALH 90h onwards reaching to 20C till LL3 (Fig. 6 A) which might be completed during the pupal stage (not evaluated). This analysis also indicates that the rate of replication accelerates after ALH 48hr with a rate of approximately 0.2 DNA content units per hour to complete the second round of replication (Fig 6A). Following replication, first complete nuclear division occur around 65-72 hours (Fig. 1F, H).

**Fig 6:**
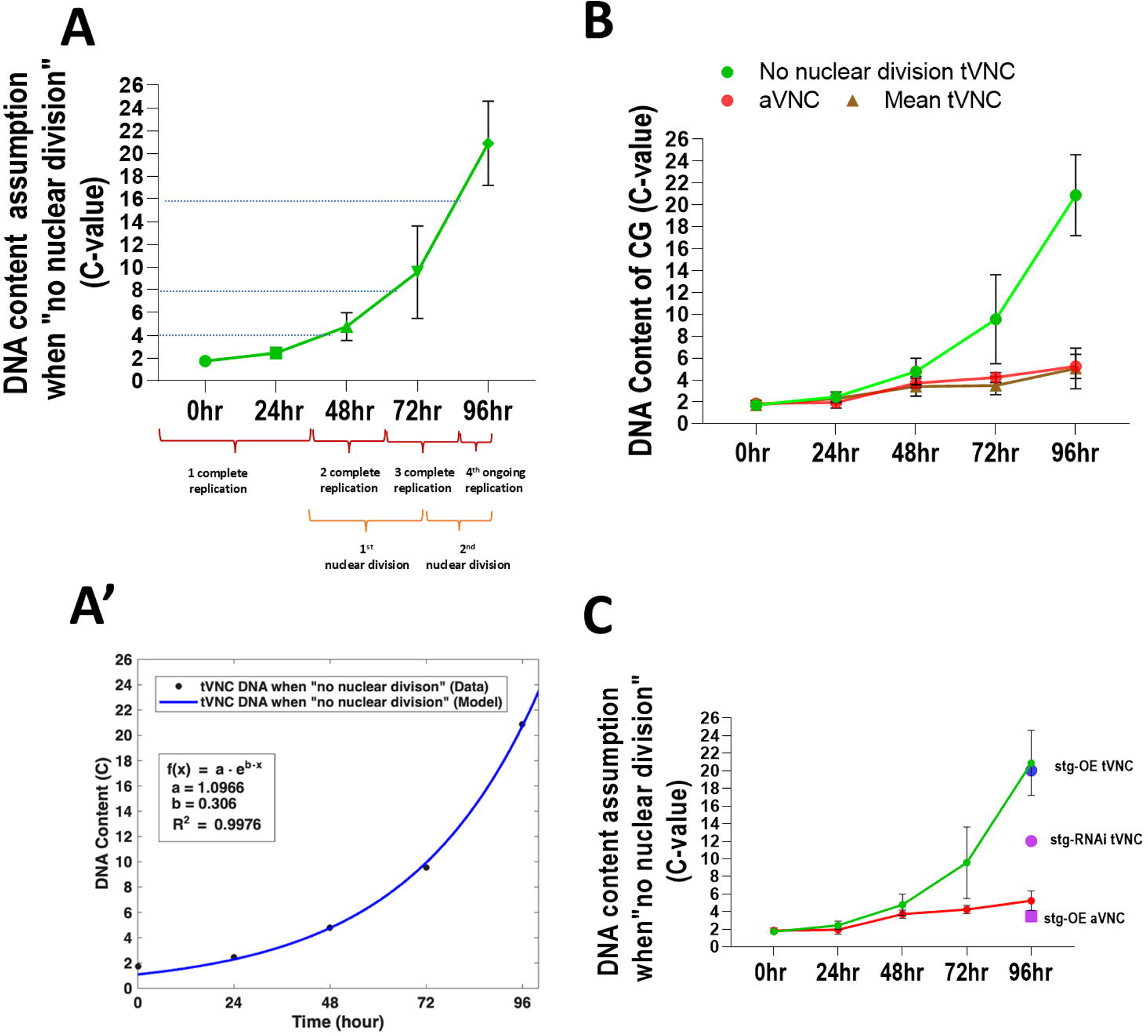
Region-specific differential regulation of the cell cycle determines the ploidy of CG nuclei. A) Quantification of DNA content of tVNC CG nuclei under “no nuclear division assumption”, showing that DNA content increases exponentially from ALH 48hr to ALH 96hr, by three complete and fourth ongoing replication rounds, A’) Mathematical model showing the same. B) The comparison of DNA content in tVNC CG nuclei, based on the “no nuclear division assumption” (green), is illustrated alongside the average DNA content of tVNC (brown) and aVNC CG (red). This shows that, under the no nuclear division assumption, DNA content increases exponentially. In contrast, the average DNA content in both tVNC and aVNC increases linearly, despite the different cell cycle dynamics observed in the two regions. C) The comparison of DNA content between tVNC and aVNC CG nuclei, under conditions of stg misregulation, is analyzed alongside DNA content in the “no nuclear division assumption” (indicated in green) and the aVNC (red) controls. In the *stg*-depleted condition, tVNC CG continues replication and reaches a 12C DNA content in LL3, entering the exponential phase; however, its replication rate does not match that of the control. Conversely, stg overexpression (stg-OE) does not impact the total DNA content. When stg is ectopically expressed in aVNC, the DNA content returns to levels between 2C and 4C.

The rate of mitosis accelerates post ALH 65-72h and is equal to 0.035 (represented by b in the Model Equation 1). Our analysis indicates that the CG in tVNC increases their nuclear count by two full rounds of nuclear division. Also, the DNA content increases by undergoing three complete rounds of replication at varying rates but maintaining a ploidy primarily between 2C and 4C per nucleus.

Next we comparedDNA content of aVNC CG nuclei, which do not divide during different developmental stages, with the calulated DNA content of tVNC under no nuclear division assumption (Fig. 6B). The mean DNA content of aVNC per nucleus increases linearly and does not enter an exponential phase at ALH 35h (Fig. 6B). Interestingly, after ALH 48h we observe a clear bifurcation in the rate of DNA increase in tVNC CG (with the exponential growth rate b value to be 0.0278) and aVNC CG (slope of the line a value is 0.0378), showing spatial differences in the replication/endocycle rate. However, the mean DNA content of both tVNC and aVNC across different developmental stages is similar, ranging from 2C to 6C. Thus, indicating that despite the differences in cell cycle in tVNC and aVNC CG both maintain the similar DNA content per nucleus.

We noted earlier that the knockdown of the M-phase regulator stg halts nuclear division in tVNC CG, causing these nuclei to switch from replication to endocycle mode and increase their DNA content per nucleus (Fig. 6C). To assess the rate of this increase compared to the control, we compared the data with our no-nuclear division assumption model. We found that the tVNC CG depleted of *stg* continues DNA replication, enters the exponential phase, but does not match the replication rate of the control (Fig. 6C). This suggests that CG in tVNC can switch from replication to endocycle mode, increasing their DNA content when nuclear division ceases. Conversely, ectopic expression of *stg* induces more nuclear division in CG, maintaining a 2-4C DNA content even during LL3, without affecting the total number of replications rounds when compared to the no-nuclear division assumption (Fig. 6C).

We also examined stg regulation in aVNC CG. Upon ectopic expression, we found that endocycling nuclei switch to mitotic mode, reducing the DNA content to 2-4C (Fig. 6C). Together, our data demonstrate that CG in tVNC and aVNC follow the normal and endocycling modes of replication, respectively; the availability of *stg* being crucial in determining the path taken.

### 7: Cell cycle regulators are necessary and sufficient to maintain the morphology of tVNC cortex glia

CG are known to form a unique reticulate network, having multiple thick and thin branches which together form a honeycomb-shaped structure, often termed as trophospongium around neural cell bodies (Hoyle, 1986). A single CG may ensheathe around 100 neuronal cell bodies (Awasaki et al., 2008). Since we know that cell cycle regulators are required for DNA replication and nuclear division in CG, we further examined whether disrupting these regulators also affects the complex morphology of these cells.

To assess CG morphology, we quantified the volume by marking the CGs with mcd8-RFP, driven by a cortex glial specific Gal4 driver, while simultaneously knocking down several cell cycle regulators in these cells. Our findings revealed that the knockdown of *dup*, *CycE*, *dacapo*, and *stg* significantly reduced the volume of tVNC CGs to 0.027 (*dup RNAi*), 0.13 (*CycE RNAi* #2), 0.10 (*CycE RNAi* #1), 0.79 (*stg-RNAi* #1), 0.78 (*stg-RNAi* #2), and 0.69 (*stg-RNAi* #3) fold (Fig. 7C, D). The CGs depleted of cell cycle regulators displayed thinner, more disrupted main branches than the control, along with a significant reduction in side branches that enwrap the progeny (Fig. 7A, B). Notably, following *CycE* knockdown, we observed an accumulation of CG membrane, a phenotype associated with blocked vesicular transport and a loss of proliferation (Coutinho-Budd et al., 2017).

**Fig 7:**
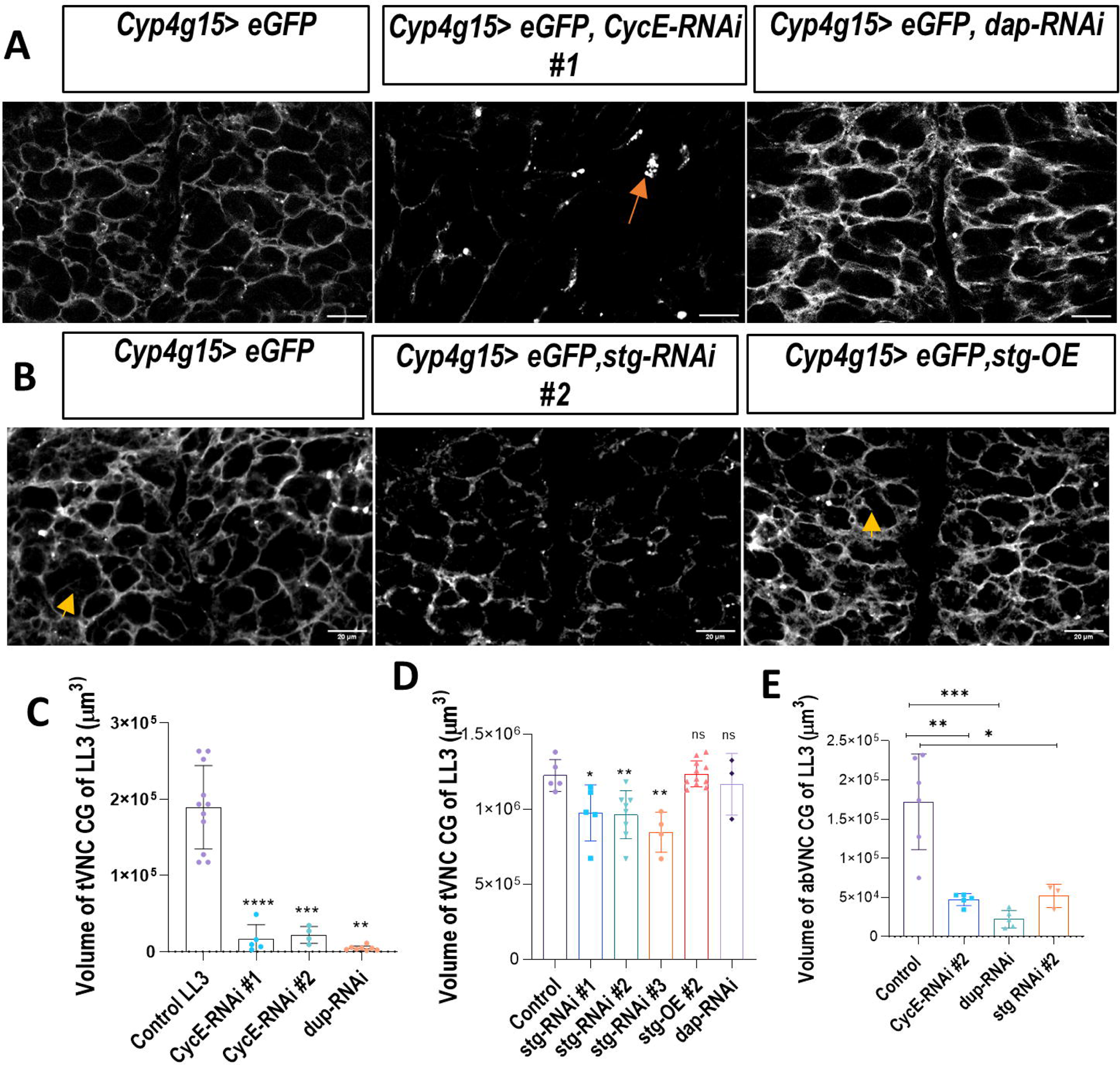
Cell cycle regulators are required for CG membrane growth. A) Projection of a few ventral sections shows that the morphology of CG in tVNC is affected in the *CycE-RNAi*, orange arrow marks the accumulation of membrane vesicles, and in the *dap-RNAi,* the main branches are thicker than in the control. B) Projection of a few ventral sections shows that the morphology of CG in tVNC upon disruption of mitotic regulator *stg,* CG chambers become disconnected, and also there is loss of finer branching, while in overexpressing *stg*, both the main and side branches become denser. C) Quantification of volume of CG in tVNC in different knockdown condition like *dup-RNAi* and *CycE-RNAi* (#1,#2) D) Quantification of volume of CG in tVNC in different conditions like *stg-RNAi, stg-OE* and *dap-RNAi* E) Quantification of the volume of CG in aVNC in different knockdown conditions, such as *dup-RNAi.CycE-RNAi*, and *stg-RNAi*, The statistical evaluation of significance based on the unpaired *t*-test is marked with asterisks, \*\*\*\**P* < 0.0001, \*\*\**P* < 0.001, and \*\**P* < 0.01. Scale bar: 20 µm

Similar to tVNC, aVNC CG also exhibits disrupted morphology when *dup*, *CycE*, and *stg* are knocked down. Although these cell cycle regulators do not directly affect the number of nuclei, their knockdown reduces the rate of replication in hyperploid cells. Additionally, the volume of aVNC CG shows a significant reduction to 0.13 (*dup-RNAi*), 0.27 (*CycE RNAi*), and 0.30 (*stg RNAi*) fold (Fig. 7E), indicating that decreased ploidy may also impact membrane growth in these CG cells. Furthermore, when we provided *CycE* and *stg* ectopically, we did not observe any significant changes in the volume of tVNC CG, however, both the main and side branches appeared thicker compared to the control (Fig. 7C, D).

Overall, our data suggest that CG morphology is associated with cell cycle regulation in both tVNC and aVNC.

## Discussion

Our study investigates the regulation of the variant cell cycle in CG of VNC region. We uncovered an intriguing pattern of CG growth in the tVNC compared to the aVNC regions. Cyclins and cyclin-dependent kinases play a crucial role in regulating CG cell cycle and growth, with replication checkpoints largely remaining inactive. Furthermore, we identified region-specific differences in CG cycling behavior between the thoracic and abdominal regions of the VNC. Our findings demonstrate that CG nuclei in the tVNC increase their DNA content and divide the nucleus, maintaining ploidy largely between 2C and 4C during most of larval development, whereas CG nuclei in the aVNC exclusively undergo endocycling. This study provides insights into the growth and cell cycle regulation of CG.

### Polyploidy and various ways of Non-Canonical Cell Cycling

Polyploidy is a conserved biological mechanism observed in various organisms, including protists, algae, angiosperms, molluscs, insects, mammals, and humans (Brodskiĭ&Uryvaeva, 1985; Nano et al., 2019; Ramesh et al., 2025). While polyploidy was once linked to an increased risk of cancer and tumor growth, it is now recognized for its vital role in numerous biological processes. These processes include managing stress responses, promoting wound healing, enhancing DNA damage resistance and repair, facilitating the formation of binucleate alveolar cells in the lactating mammary gland, supporting the development of megakaryocytes, and many others (Cohen et al., 2021; Fox &Duronio, 2013; Øvrebø& Edgar, 2018; Zielke et al., 2013).

Polyploidy can result from changes in the cell cycle, which involves several processes (Edgar et al., 2014; Eikenes et al., 2015; Fox &Duronio, 2013; Frawley & Orr-Weaver, 2015; Zielke et al., 2013). These processes include endocycling, endoreplication, or endoreduplication, where only the G/S phases occur without any M phase. Additionally, two more forms of the cell cycle can be observed based on the completeness of cytokinesis. The first is endomitosis, where cytokinesis does not occur. The second is acytokinetic mitosis, which leads to the formation of compartments within a cell due to incomplete cytokinesis (Edgar et al., 2014; Eikenes et al., 2015; Fox &Duronio, 2013; Frawley & Orr-Weaver, 2015; Zielke et al., 2013).

A combination of various non-canonical cell cycles can operate together in a cell, as seen in both *Drosophila* and mammalian systems (Cohen et al., 2021). For instance, in *Drosophila*, follicle cells in the ovaries and accessory glands in the testes initially undergo mitosis before switching to endocycles (Box et al., 2019; Deng et al., 2001; Shcherbata et al., 2004; Sun & Deng, 2005). In contrast, the rectal papilla cells in the adult *Drosophila* gut undergo several rounds of endocycling before experiencing one round of error-prone mitosis (Duncan et al., 2010; Fox et al., 2010; Schoenfelder et al., 2014).

Similarly, in mammalian systems, there are examples of polyploid cells, such as megakaryocytes, trophoblasts, and the glandular epithelium of the endometrium, which undergo multiple rounds of replication and nuclear division (and possibly endomitosis) (Bergmann et al., 2015; Brodsky et al., 1992; Brown et al., 1997; Hesse et al., 2012; Lordier et al., 2008; Losick et al., 2013; Odell et al., 1976; Zybina et al., 2001). These examples suggest that variant cycling processes enable certain cells, under specific conditions, to increase in size or output without requiring complete cell division.

The comparative analysis of SPG and CG reveals distinct yet overlapping features in their developmental patterns and nuclear characteristics. SPG engage in repetitive cycles of endocycling alongside nuclear division, suggesting a coordinated approach to cellular growth and proliferation. Notably, SPG, particularly in the optic lobes, displays a marked increase in both the number of nuclei and DNA content, often showing polyploid and syncytial cells. In contrast, while SPG in the VNC retains polyploid characteristics, they are strictly mononucleate (Unhavaithaya & Orr-Weaver, 2012; Von Stetina et al., 2018). In CG, multiple rounds of replication and nuclear division take place to form a dense reticulate network. The growth pattern of CG is largely known of those present in VNC and they are known to have high DNA content(Rujano et al., 2022; Yadav et al., 2024). Here we show that individual nuclei in CG network present in tVNC maintain their ploidy between 2C-4C. These CG in tVNC show a pronounced spatial distribution in the VNC, with exponential increases in nuclei number during larval development. On the other hand, those present in abdominal region primarily undergo endocycling, resulting in mononucleate polyploid cells. These variations highlight not only the functional similarities in cyclic growth patterns between SPG and CG but also underscore the importance of regional cues in dictating their morphological and ploidy differences. Further investigation warranted to elucidate detailed regulatory mechanisms involved in the formation of complex glial niche around neural cells.

### Importance of cell cycle regulators and mitotic checkpoints in controlling variant cell cycle

Endocycles produce an increased nuclear size and content within a single nucleus (ex, *Drosophila* salivary gland, rectal papillae cells, and trophoblast giant cells in mammals). In endomitosis and acytokinetic mitosis, failure or incomplete cytokinesis results in the production of binucleate/multinucleate polyploid cells (e.g., cardiomyocytes and hepatocytes). Thus, to make a syncytial polyploid cell, nuclear division in an endocycling cell is necessary. It is interesting to note that in megakaryocytes which usually undergo endomitosis, loss of CDK1 results in a switch in their cycling mode from endomitosis to the endocycle, allowing them to remain polyploid and continue functioning normally (Trakala et al., 2015). Similarly, stg/cdc25, an essential regulator of mitosis, determines the nuclear division in polyploid SPG and CG in the *Drosophila* nervous system (Beachum et al., 2025; Von Stetina et al., 2018). Knockdown of stg prevents multinucleated SPG from undergoing endomitosis, while their overexpression makes even the mononucleated SPG undergo nuclear division, including those present in VNC (Von Stetina et al., 2018). This spatially restricted regulation of nuclear division is very interesting. The same phenomenon we have observed in the case of CG, where the hyperdiploid CG undergoes nuclear division upon stg overexpression in the abdominal region of the VNC. Understanding the regulatory mechanisms that govern the balance between endocycle and nuclear division, as well as identifying the specific cell-autonomous and non-cell-autonomous triggers for these processes, is a fascinating and crucial area for further study.

Normal DNA replication during the typical cell cycle is regulated by various cell cycle regulators and checkpoints, ensuring accurate genome duplication. In certain cell types that undergo endocycling, continuous replication occurs without the active involvement of replication checkpoints, which helps prevent p53-mediated apoptosis caused by checkpoint activation (Lilly and Spradling., 1996; Mehrotra et al.,2000; Zhang et al., 2014). In cases where polyploid mitosis is observed, such as in SPG and rectal papilla cells, these cells exhibit error-prone mitosis. This is characterized by the formation of chromosome bridges, lagging chromosomes, chromosome fragments, and delayed mitotic transitions (Fox et al., 2010; Schoenfelder et al., 2014; Unhavaithaya & Orr-Weaver, 2012). CG nuclei have also been reported to display variable chromosome numbers (Rujano et al., 2022) during larval development. We demonstrate that CG in VNC undergoes multiple rounds of both endocycling and mitosis, regulated by canonical cell cycle regulators, but without the active involvement of most replication checkpoints. This modification likely arises from the continuous need to increase DNA content for cellular growth to accommodate neural cells. Future research will help us understand the implications of this modification and the specific replication strategy in these cells.

Our work demonstrates that CG is a unique model for studying variant cell cycle mechanisms in the nervous system. It also underscores the importance of further investigation into cell-intrinsic and extrinsic molecular pathways, which could provide valuable insights into both normal development and disease states in which ploidy is altered, such as in glioma or other cancers.

## Methods

### Fly rearing and handling

*Drosophila* melanogaster was maintained at 25°C ±1, and RNA interference (RNAi) crosses were conducted at 29°C using a standard nutritional medium comprising sugar, agar, maize powder, and yeast. Appropriate fly crosses were established following standard protocols to produce progeny with the desired genotypes (Yadav et al.,2024). The following fly stocks were utilized in the experiments: Oregon R+ as the wild-type, *Repo-GAL4* (BL 7415), *Cyp4g15-GAL4* (39103), *UAS-eGFP*(II) (BL-5431), mCD8RFP (BL-27398), and Fly-FUCCI (BL-55121), Stg-RNAi (BL-36094,34831, V17760*), stg-OE* (BL-4778,58439), *CycE-RNAi* (29314, v10204), *CycE-OE*(BL-4781), and *dup-RNAi* (BL-29562), *grp-RNAi*(BL-36685), *tefu-RNAi*(BL-31635),, *loki-RNAi*(BL-35152), *ATM* (31635),, *ATR* (VDRC 103624),, *Chk1-OE* (Kizhedathuet al 2021), Chk1-RNAi(V110076), cdk1-RNAi(VDRC 106130), dacapo-RNAi(bl-36720,64026), *dacapo-OE* (bl-83334), *cdc-6-RNAi*(BL-55734), *gem-RNAi* (bl-30929,) *gem-OE*(bl-53720), *MCM10-RNAi* (VDRC 9241).The following genotype combinations were generated in the laboratory for the study: *UAS-eGFP; Cyp4g15-GAL4, UAS-FUCCI; Cyp4g15-GAL4, mCD8RFP; Cyp4g15-GAL4, Stringer-red; Cyp4g15-GAL4*

### Immunostaining, confocal microscopy, and documentation

Larvae at specific developmental stages, including early first instar (ALH 0-3hrs), L2 (ALH 24-28hrs), early third instar (ALH 48-54hrs), mid third instar (ALH 72hrs), third instar (ALH 93-96hrs), and wandering larvae as late third instar (referred to as LL3 throughout the text), were selected from the F1 progeny of respective crosses. The central nervous systems (CNS) were dissected in Phosphate Buffer (PBS 1X containing NaCl, KCl, Na2HPO4, KH2PO4, pH 7.4), fixed in 4% Paraformaldehyde for 30 minutes, rinsed in 0.1% PBST (1XPBS, 0.1% TritonX-100), and incubated in a blocking solution (0.1% TritonX-100, 0.1% BSA, 10% FCS, 0.1% deoxycholate, 0.02% thiomersal) for 30 minutes on a rotator at room temperature. The samples were then incubated with the necessary primary antibody at four degrees Celsius overnight. The antibodies used include mouse Anti-Repo (1:100, Developmental Studies Hybridoma Bank, 8D12), and rabbit anti-GFP antibody (1:200, Invitrogen, A-11122). The following day, samples were washed three times in 0.3% PBST and incubated with the appropriate secondary antibody at a 1:200 dilution either overnight or for two hours at room temperature. The secondary antibodies used were donkey anti-rat Alexa 488 (Invitrogen, A-21208), Goat anti-Mouse Alexa Fluor 568 (cat. No. A11004), and Chicken Anti-Rabbit Alexa Fluor 488 (Invitrogen, A-21441). After incubation with secondary antibodies, the samples were washed three times in 0.1% PBST and counterstained with DAPI (1µg/ml, Invitrogen, D1306) at 4°C overnight when necessary. Following the final washes in 0.1% PBST, the samples were mounted in DABCO (Sigma, D27802) for further analysis. Images were captured using the following confocal microscopes: Zeiss LSM-510 Meta at the Department of Zoology, and Leica SP8 STED facility at CDC, BHU. All images were quantified using Fiji/ImageJ software (NIH, USA) and assembled using Adobe Photoshop and MS PowerPoint software.

### Image analysis using Fiji/ImageJ software

Confocal imaging of CG in the tVNC and aVNC was examined using Fiji/ImageJ software (NIH, USA) to assess the number, area, and volume. The freehand selection tool was employed to manually delineate the region of interest (ROI) for integrated intensity measurements. The multipoint selection tool was used to manually count the number of glia nuclei (GFP+, Repo+), based on GAL4-driven GFP and Repo staining. To determine the size of CG nuclei (GFP+, Repo+), the area was outlined using the freehand selection tool around Repo staining, followed by the analyze-measure option.

Integrated density for DAPI in the CG nuclei was measured by manually outlining each GFP+, Repo+ DAPI+ nucleus, with the area outline done using the freehand selection tool around Repo staining. The confocal scanning parameters for each experimental setup were kept constant for all intensity quantifications (Yadav et al., 2024). Z-stacks that covered the entire nucleus were acquired. Controls were analyzed alongside each experimental set. We measured around 20% CG nuclei in the ventral sections, leaving the CG in the dorso-lateral and neuropile regions due to overlapping neural progenies.

DNA content was quantified by obtaining the sum of the DAPI integrated density in Image J and then the mean/quartiles-integrated density of CG was normalised with that of sperm (1C).

### Mathematical modelling

#### I. Exponential Growth Kinetics of Nucleus and DNA in the Thoracic Cortex Glial Cells of *Drosophila*

We model the temporal increase in mean nuclei values and the total DNA content (estimated) of thoracic cortex glia using an exponential growth function. The mathematical equation used for both variables is as follows.

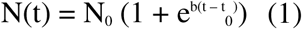

Here, N(t) represents the mean nuclei count at time t, N_0_ represents the initial mean nuclei count, b is the exponential growth rate, and t_0_ represents the transition time at which rapid growth occurs. We used MATLAB’s Curve Fitting Toolbox to fit the data with the numerical solution of Equation (1). The fitted curve closely matches the data points showing high goodness of fit (R² = 0.9902).

We estimated the total DNA content, using the relation between mean nuclei value and mean DNA intensity as follows.

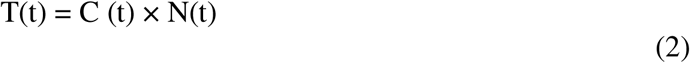

where C(t) represents mean DNA content per nucleus at time t, and T(t) represents the total DNA content at the time t, respectively. The resulting curve also showed high goodness of fit (R² = 0.9834) with the data suggesting exponential increase in the total DNA content over time.

##### No nuclear division assumption

We computed how the DNA content would vary if there were no nuclear division in tVNC CG. The mean number of nuclei increased from ALH 0h to ALH 96h, we assume that the number of nuclei remains constant throughout the larval stage at 63.67, the number observed during ALH 0h i.e. in this period of development, no nuclear division has taken place. We divided the total DNA content T (t) with the mean nuclei number N_0_ at ALH 0 hr when no nuclear has taken place. We applied the same method at each developmental stage to estimate DNA content when no nuclear division has occurred.

### Volume calculation

For volume analysis of CG, a default thresholding was performed to select pixels of interest based on the RFP intensity of the pixel values. Thereafter, the measure stack plugin was used to find the fluorescent area (RFP) of each CG section of tVNC and aVNC through the analyse-measure option. The area obtained was thus multiplied by the number of stack intervals (1) to determine the volume of CG.

Statistical analysis was performed using GraphPad Prism 9 software, where a two-tailed unpaired t-test was used to assess the statistical significance between the mean values of the two groups, with asterisks indicating significance as follows: n with p < 0.05 (**** if p<0.0001, *** if p<0.001, ** if p<0.01, and * if p<0.05) considered statistically significant, and if p>0.05 considered non-significant.

## Acknowledgement

We are grateful to Profs. S C Lakhotia, R. Raman, P. Sinha, and G Pandey, for critically reviewing the manuscript, insightful comments, and valuable suggestions throughout. We also thank Prof. Calvi, Dr. Krishanu Ray for generously sharing the reagents and fly stocks. We also thank the Bloomington *Drosophila* Stock Center (BDSC) for providing fly stocks. The national facility of Zeiss 510 Meta and Zeiss LSM 900 confocal microscopy at the Department of Zoology, and Zeiss 510 Meta at ISLS BHU, Leica SP8 STED confocal microscopy facility at CDC, BHU are also acknowledged. Finally, we thank the Cytogenetics Laboratory, Department of Zoology, for providing us with all the necessary instrumentation facilities.

## Funding

This work was supported by grants from the DST-SERB (ECR/2018/002837, CRG/2022/006350), CST-UP (5653), and BHU-IoE to R.A., and University Grants Commission, India, for startup grant and BHU-IoE to R.P.,BHU Non-NET-RET and CST-UP (5653), fellowship to V.Y., and Council Of Scientific And Industrial Research (CSIR), India for providing NET-JRF to R.K.M, Prime Minister Research Fellowship (PMRF), from the Ministry of Education, Government of India, to S.T.

## Conflict of interest

The author (s) declare no conflict of interest.

## Data Availability

Fly lines are obtained from Bloomington Stock Center, USA, and genetic combinations created are available upon request. The authors affirm that all other data necessary for confirming the conclusions of the article are present within the article, figures, and tables.

---

**Fig S2: Validating *CycE* and *dap* regulation in CG cell cycle with different Fly lines**

A) Control (*Cyp4g15>eGFP*) showing 3-4 CG nuclei per hemisegment in ventral section projections, making a patterned arrangement, at ALH 24hr *CycE* misexpression CG shows no significant change in nuclei number,

B) Knockdown of *CycE* with the second RNAi line and its ectopic expression with the second line resulted in a decrease and an increase of CG nuclei number, respectively, as seen in Fig. 2.

C) At the LL3 stage, *dacapo* knockdown with the second RNAi line also shows a similar result, increasing the number of nuclei in comparison to the control.

## Notes

### Competing Interest Statement

The authors have declared no competing interest.

